# High Concentrations of floating life in the North Pacific Garbage Patch

**DOI:** 10.1101/2022.04.26.489631

**Authors:** Fiona Chong, Matthew Spencer, Nikolai Maximenko, Jan Hafner, Andrew McWhirter, Rebecca R. Helm

## Abstract

Floating life (obligate neuston) is a core component of the ocean surface food web. However, only one region of high neustonic abundance is known so far, the Sargasso Sea in the Subtropical North Atlantic, where floating life provides critical habitat structure and ecosystem services. Here, we hypothesize that floating life is also concentrated in other gyres with converging surface currents. To test this hypothesis, we collected samples through the eastern North Pacific Subtropical Gyre in the area of the North Pacific “garbage patch” (NPGP) known to accumulate floating anthropogenic debris. We found that densities of floating life were significantly higher inside the central part of NPGP than on its periphery, and there was a significant positive relationship between neuston abundance and plastic abundance. This work has important implications for the ecology and human impact of subtropical oceanic gyre ecosystems.

## Introduction

Marine surface-dwelling organisms (obligate neuston) are a critical ecological link between diverse ecosystems [1]. Hundreds of species that live in the water column, seafloor, or even in freshwater spend part of their lifecycle at the ocean’s surface (see review in [1]). As floating organisms, obligate neuston are transported and concentrated by ocean surface currents.

Many species of neuston are globally distributed, but currently only one ocean region is known to concentrate neuston into high densities. The Sargasso Sea is named for the neustonic *Sargassum* algae and is a marine biodiversity hotspot supported by neuston. The Sargasso Sea is critical to the ecology of the North Atlantic and provides millions to billions of US dollars in ecosystem services annually [2,3]. But is the Sargasso Sea the only region of the world’s oceans where floating life concentrates?

Plastic pollution, transported by the same surface currents that transport neuston, provides a clue: large amounts of floating debris are transported to and concentrated in “garbage patches” identified in all five main subtropical gyres of the North Atlantic (the Sargasso Sea), South Atlantic, Indian Ocean, North Pacific, and South Pacific [4,5]. Neuston, subjected to the same oceanographic forces that move buoyant man-made waste and pollutants, may also be concentrated in ‘garbage patches’. We hypothesize that these regions could be neuston seas, like the Sargasso Sea, and could provide similarly critical ecological and economic roles.

Convergence of neuston life into high densities may be critical for many neustonic species, and the organisms that depend on them. Many neuston, including foundational members of the neuston food web, *Physalia, Velella,* and *Porpita,* are incapable of swimming or directional movement. Predatory neuston such as the blue sea dragon *Glaucus* and the violet snails *Janthina* also lack the ability to direct their movement, and must physically bump into prey in order to feed [6,7]. Even more strikingly, *Glaucus* and some species of *Janthina* must also be in physical contact to mate [8–10]. These adaptations point to the need for extremely high-density regions in order for these species to survive and reproduce. Some members of the neustonic community may also have adaptations to survive in relatively low nutrient waters (characteristic of many subtropical gyre regions [11]), including the presence of endosymbiotic zooxanthellae [12], similar to those found in corals (e.g. *Velella* and *Porpita;* Figure 1). Neuston are in turn consumed by fish, seabirds, and turtles [1], which may seek out dense concentrations as feeding grounds. Over 100 species of fish and diverse invertebrates [13–15] also live at the surface and may shelter with, feed upon, and be eaten by obligate neuston.

**Figure 1.**
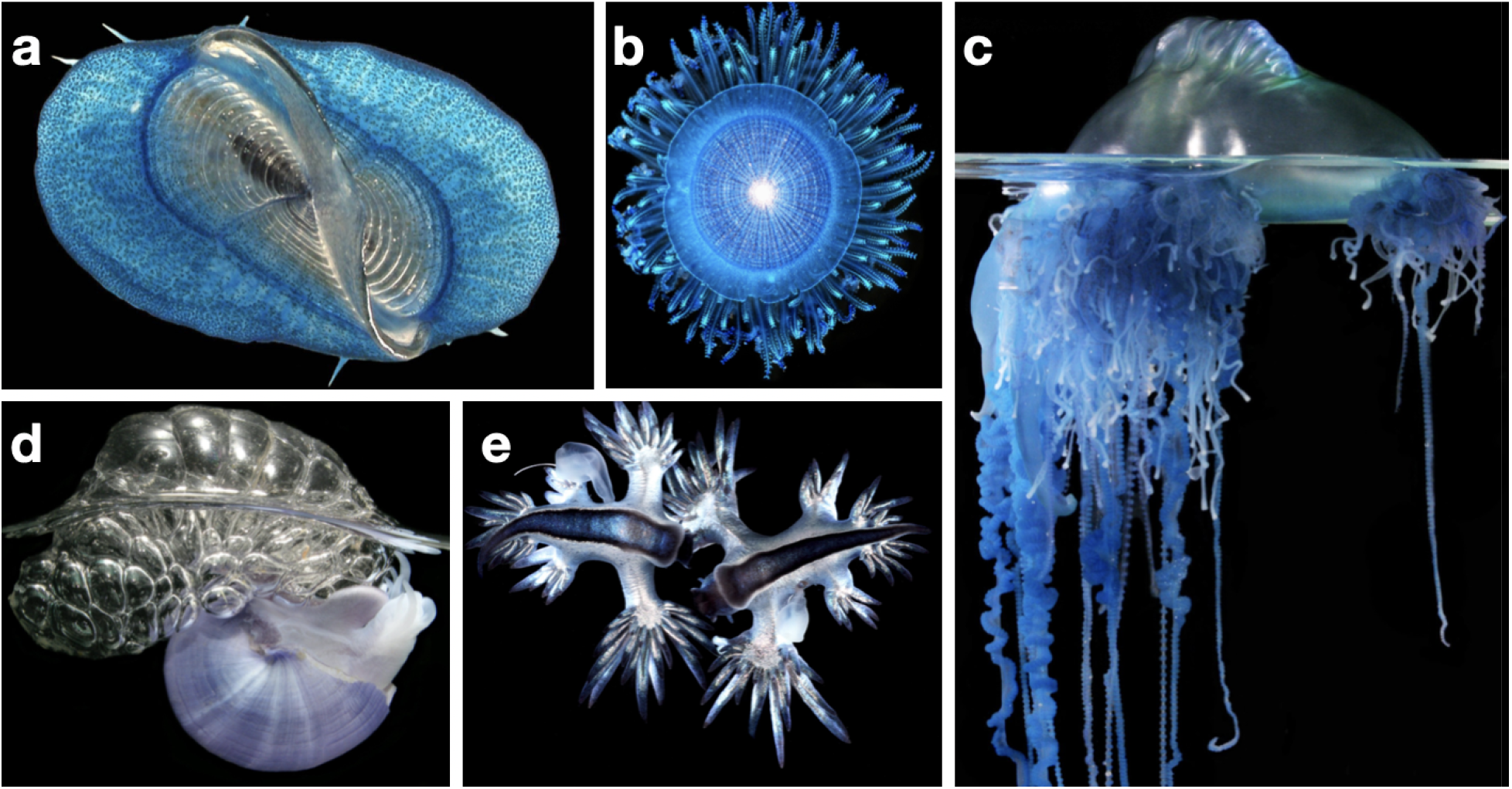
The neustonic organisms represented in this study, based on Helm 2021. (a) top-down view of by-the-wind sailor *Velella* sp. (b) top-down view of blue button *Porpita* sp. (c) side view of Portuguese man-o-war *Physalia* sp., (d) side view of violet snail *Janthina* sp. (e) top-down view of the blue sea dragons *Glaucus* sp. Images by Denis Riek.

The North Pacific Garbage Patch (NPGP) is the largest and most infamous of the garbage patches [16]. It exists within the North Pacific Subtropical Gyre (NPSG), a massive semi-enclosed region characterized in part by comparatively low nutrient densities [17,18]. Diverse neustonic species are documented from the NPSG [19–21], including several species of blue sea dragons *(Glaucus* spp.), that has been found nowhere else [20]. Because the NPGP is thousands of miles from shore, has a dynamic spatial structure, and exhibits significant variations in time, few surveys of neuston have been performed in the NPGP.

To test our hypothesis that garbage patches may also be neuston seas, including the NPGP, we conducted a community science survey through the NPGP with the sailing crew accompanying long-distance swimmer Benoît Lecomte (https://benlecomte.com/) as he swam through the NPGP (The Vortex Swim). The sampling scheme was coordinated with our model predicting the areas of high densities of floating objects. We found increased concentrations of floating life in the NPGP and a significant positive relationship between the abundance of floating life and floating plastic. Further, neuston densities in the NPGP are among the highest ever described. “Garbage Patches” may be overlooked areas of high neuston abundance, and could serve similar ecological roles to the North Atlantic Sargasso Sea, providing food and habitat for diverse species and valuable economic services. There is an urgent need to better understand these ecosystems and the impact of plastic debris.

## Methods

Neuston samples were collected by The Vortex Swim, an 80-day sailing expedition through the NPGP. A numerical drift model was used to plan the route of the expedition in accordance with regions of predicted high concentrations of floating plastic marine debris (Fig. 1). This model has been successfully used previously to simulate trans-Pacific drift of debris generated by the 2011 tsunami in Japan [22].

### Sampling

The Vortex Swim expedition aboard the sailing vessel *I Am Ocean* started in June 2019 from Honolulu, Hawaii and reached San Francisco, California in August 2019. During this 80-day expedition, as part of a community science initiative, 22 samples were taken, with 12 in the central region of the NPGP, and 10 peripheral to or outside of the NPGP (Figure 2, Supplementary ‘site_information.xlsx’). Either a Manta trawl (width x height: 0.9 m x 0.15 m) or a neuston net (width x height: 1 m x 0.25 m) was towed along the sea surface for 30 minutes at each site (Supplementary ‘site_information.xlsx’), with an attached General Oceanics Mechanical Flowmeter (model 2030R) to measure the approximate water volume filtered by the trawl (with the exception of TLS_101 and TLS_105: no volume was recorded so volumes for these samples were manually calculated using distance traveled; see site_information.xlsx for more information).

**Figure 2:**
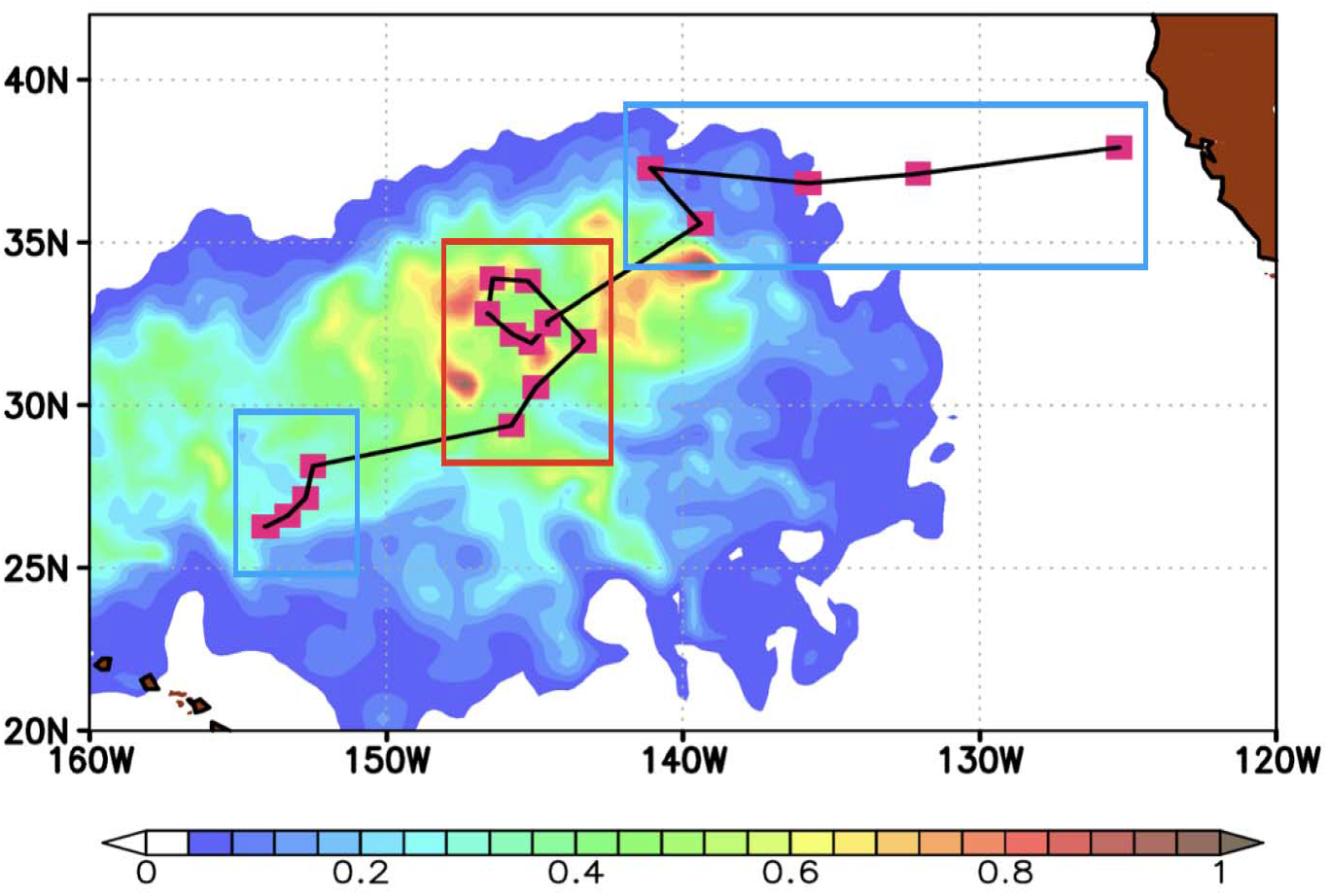
Sampling stations and model tracer concentration, outlining the NPGP pattern during The Vortex Swim expedition. Higher predicted floating particle concentrations are red, lower are blue and white, units are relative to the maximum concentration. Samples outside of the garbage patch (or NPGP) are boxed in blue, samples within in red.

Due to the fragility of neuston and difficulty of sampling, biological preservation was not possible and we used a photographic survey for our analysis (Figure S1). One image was taken per sample, with the exception of SJR_019, where two images were taken as organisms and plastic in the sample were much more abundant (refer to supplementary ‘averages_no_rounding.csv’). All neustonic organisms, plastic, and other inorganic particles present on each image were identified, counted, and recorded using JMicroVision v1.3.2 (Roduit, 2019). Nothing below approximately 0.5 mm in the longest dimension was counted. Counts were standardized to numbers per cubic meter (referred to as density throughout this text). Organisms were identified to the lowest taxonomic level possible: for all true neuston this was to the genus level. Obligate (true) neuston counted here consist of *Velella, Porpita, Janthina, Glaucus,* and *Physalia* only (Table 1).

**Table 1.** Taxa observed from the image analysis of Vortex Swim manta trawl or neuston net samples.

| True neuston taxa | Other taxa | Inorganic objects |
| --- | --- | --- |
| *Velella* | *Halobates* | Wood |
| *Porpita* | *Planes* | Rock |
| *Janthina* | Copepods | Rope |
| *Glaucus* | Unidentified blue dots | Plastic |
| *Physalia* | Fish |  |
|  | Nudibranch |  |
|  | Gastropod |  |
|  | Larvae |  |
|  | Ostracod |  |
|  | Isopod |  |
|  | Unknown crustacean |  |
|  | Unknown gelatinous organisms |  |

### Model tracer simulations

Accumulation of marine debris and neuston in the garbage patch was simulated in numerical experiments using velocities from the Surface Currents from Diagnostic (SCUD) model [23].

These velocities are derived from the historical dataset of drifter trajectories collected by the Global Drifter Program (https://www.aoml.noaa.gov/phod/gdp/) and include geostrophic currents, calculated from satellite altimetry, and wind-driven currents regressed to the local wind measured by satellite scatterometers (QuikSCAT & ASCAT). The use of Lagrangian data warrants adequate representation of the complex wind effects, combining turbulent mixing, Ekman currents, and Stokes drift due to wind waves.

The majority of anthropogenic debris originates from land-based sources whose intensities and locations are not well documented. The influence of these uncertainties of the source on debris patterns is small in the garbage patches where debris items reside for a long time (e.g., [24]), during which they “forget” their origin. To simulate the garbage patch, a constant (in time and intensity) tracer flux was set up from all coastal grid points, and the model was looped between years 1992 and 2020 under a weak dissipation, representing degradation of debris due to physical factors (UV and storms) and biological interactions [25] until model solution saturated to 95% (the root-mean-square difference between subsequent cycles). The tracer concentration map is shown in Figure 2.

### Processing and statistical analysis

To reduce the effects of complex algorithms on our results, we scaled trawl counts with the water volume measured directly by a flowmeter and avoided conversion into counts per unit area, commonly used in “microplastic” studies and requiring additional, often inaccurate, adjustments to the wind.

Analyses of plastic and neuston concentrations were run in R 4.1.0 [26]. We used linear regression to determine whether a relationship exists between plastic and neuston abundance in the NPGP. Sites in the red box of Figure 1 were considered to be ‘in’ the NPGP, while those in the blue boxes were considered to be ‘out’ of the NPGP. Kruskal-Wallis tests were used to see if there were differences in plastic and neuston densities between sites in and out of the NPGP.

## Results

Median neuston densities in the central NPGP were 0.23 (interquartile range (IQR) = 0.31) neustonic organisms/m^3^, with densities reaching up to 4.89 individuals/m^3^, and were systematically higher than densities peripheral to the GPGP (0.02 individuals/m^3^; IQR = 0.03 individuals/m^3^) (Kruskal-Wallis chi-squared = 15.1, df = 1, *P* < 0.001; Figure 3). Neuston densities within the NPGP are extremely high, equivalent to the highest densities of *Velella* and *Porpita* observed by Savilov in the cross-Pacific survey [19].

**Figure 3.**
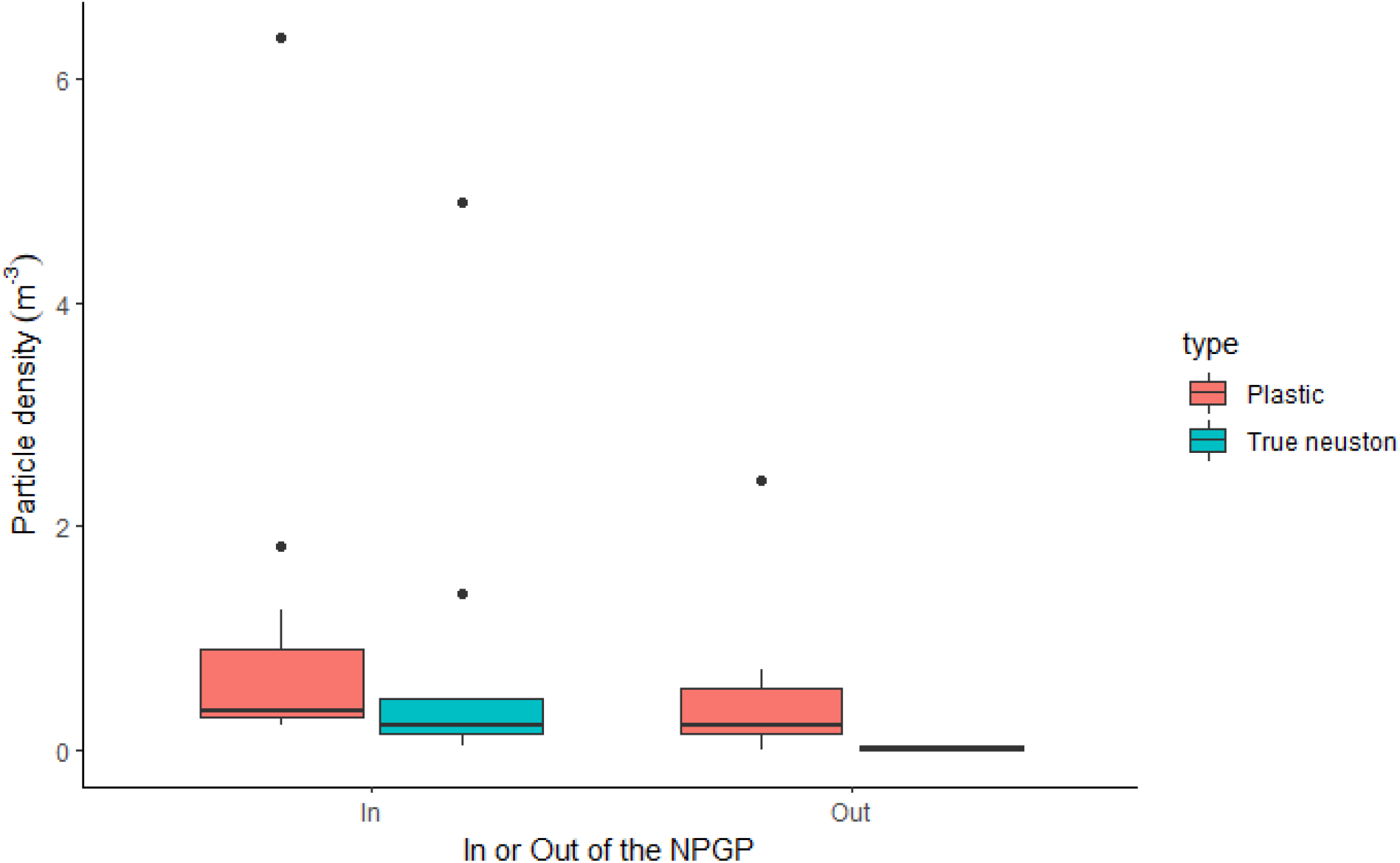
Boxplots showing the plastic and true neuston density per m^3^ sampled, within the NPGP and peripheral of it. The horizontal line is the median, the bottom and the top of the box are first and third quartiles respectively, whiskers extend to the furthest points that are no more than 1.5 times the interquartile range (IQR) from the ends of the box, and dots are potential outliers.

Median plastic densities were also systematically higher in the central NPGP (0.35 particles/m^3^; IQR = 0.60) than in peripheral (0.23 particles/m^3^; IQR = 0.40 particles/m^3^) GPGP (Kruskal-Wallis chi-squared = 4.18, df = 1, *P* = 0.040; Figure 4).

**Figure 4.**
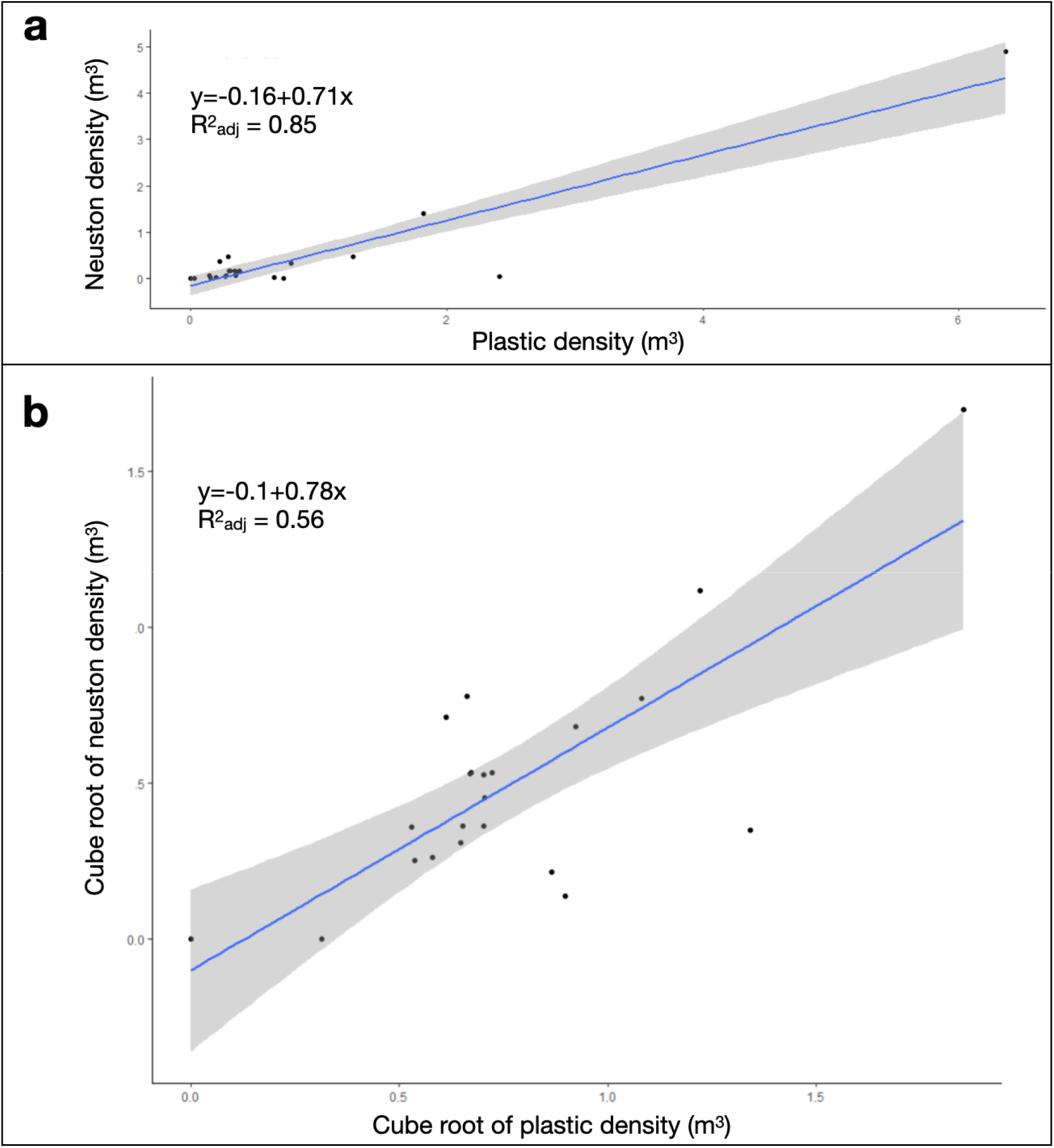
Relationship between neuston and plastic density. Black points represent sites visited by the Vortex Swim expedition. Blueline: linear regression. Gray region: 95% confidence band. The regression equation and adjusted R^2^ are also displayed. a) relationship between neuston and plastic density, b) relationship between the cube-root transformation of neuston density and the cube-root transformation of plastic density.

Using linear regressions, we found a significant positive relationship between neuston and plastic densities in the NPGP (Figure 4). In areas of high plastic density there was also high neuston density (F = 118.8, df = 1, 20, *P* < 0.001, adjusted R^2^ = 0.85). However, it is worth noting that SJR_019, which was taken within a slick, was an influential observation, with a Cook’s distance larger than 1 (Figure S2). When this observation was removed, there was still a significant positive relationship between neuston and plastic densities, although the proportion of variance explained was lower (F = 6.0, df = 1, 19, *P* = 0.02, adjusted R^2^ = 0.19; Figure S3). To reduce the influence of SJR_019, we performed a cube-root transformation of neuston and plastic density, and the relationship between plastic and neuston remained significant (F = 28.27, df = 1,20, *P* < 0.001, adjusted R^2^ = 0.56).

## Discussion

We found high concentrations of floating life and plastic in the NPGP, consistent with our hypothesis that this region may be a neuston sea. A limited number of studies have examined neuston in this region, and it is difficult to infer processes and patterns by comparing between them. Neuston concentrations in this region may vary seasonally or annually. In our study, results can be used to better understand horizontal transport of neuston, and neuston and plastic both appear to be concentrated by similar physical forces. Surface features like slicks that further concentrate neuston, may be important for neuston life history, and we found evidence that neustonic animals are reproducing in the NPGP. The possible overlap between garbage patches and neuston seas has important implications for their role in the broader ocean and potential need for protection.

The neustonic ecosystem is poorly understood, and a limited amount of work has been done on neuston and plastic distributions in the NPGP [21], however, differences in study design make it difficult to compare results across studies. For example, Egger et al. (2021) compared neuston in and around the NPGP to waters 40°N. They found key neuston species only within and around the NPGP (including *Porpita porpita, Janthina janthina,* and *Glaucus* spp.), and in our study these species are most abundant in the central NPGP. In contrast to our finding of a positive relationship between neuston and plastic densities, Egger et al. (2021) found (from 54 trawls) correlations that were close to zero between log neuston taxon densities and log plastic density, except for a moderately strong negative correlation between log *V. velella* density (which were mostly found in waters 40°N) and log plastic density. We performed a regression analysis of plastic and neuston, rather than a correlation analysis, within our study because locations were not selected to sample in an unbiased way from the populations of neuston and plastic densities. It is not clear whether Egger et al. (2021) obtained unbiased estimates of the population correlations between neuston and plastic densities because they do not provide information on how sample locations were selected. In addition, both studies have a small sample size, and understanding the relationship between plastic and neuston will require additional sampling.

Our study suggests that, when present, neuston and plastic are concentrated by similar physical forces, but unlike plastic, neuston abundance likely varies with organismal lifecycles. Our study found high total neuston numbers compared to Eggar et al (2021). For 30 minute trawls, Egger et al. 2021 found on average, 2.7 true neuston and 619 plastic particles; while our study found 202 true neuston and 291 plastic particles on average. Egger et al (2021) collected two-fold more plastic than us, while we collected 75 times more neuston. A two-fold difference in plastic is not necessarily surprising given the different sampling methods, but the difference in neuston abundance is remarkable. This difference may be due to seasonal, and possibly annual, ecosystem dynamics. A strong chlorophyll bloom (e.g., [27]) occurred in the NPGP during the Vortex Swim expedition, though it is unclear if and how blooms impact neuston population dynamics. Some samples from Egger et al 2021 were also collected in the winter, when neuston dynamics may be different. Undoubtedly, future studies should be prepared for large variations and examine seasonal and interannual dynamics.

Our sample location within the NPGP allows us to focus on the effect of currents on the movement and patterns of plastic and neuston and neglect other environmental factors (such as nutrient-rich high latitudes, as in case of Egger et al. (2021)) or anthropogenic variability (such as coastal sources of plastic). This allows us to focus specifically on the horizontal transport of neuston and plastic. As a result, we can disregard the complexity of multiscale ocean currents and the effects of vertical mixing by the wind. Figures 4 and S3 show that similar regression laws describe the correspondence between densities of neuston and plastic both on the scale of NPGP (hundreds of km) and in the ocean slicks (tens of meters). As expected [28], counts of plastic and neuston were both lower under stronger winds. The reduction of plastic at the surface under high winds was somewhat stronger than for neuston, possibly, due to a smaller size and lower buoyancy of particles (Figure S3).

Within our study, a patchy distribution of neuston and plastic at the surface may be due to small-scale (sub-mesoscale) surface dynamics like slicks, and may be important for neuston predation and reproduction. We found the highest concentration of both neuston and plastic in a slick (SJR_019), and this is true for other studies as well. For example, off the coast of the island of Hawai’i, nearly 40% of surface-associated larval fish, 26% of surface invertebrates, and 95.7% of plastic were found in surface slicks, which represented only 8% of the sea surface area of the West Hawai□i study region. [13]. In the North Atlantic, neustonic *Sargassum* is often concentrated in slicks under appropriate conditions [29,30]. Sea surface slicks create a relatively small area where diverse species come into regular physical contact through drifting. Because neustonic predators like *Janthina* and *Glaucus,* both found in our study, rely on physically contacting prey [6,7,31,32], and similarly *Glaucus spp.* and some members of the genus *Janthina* depend on direct physical contact to mate [8–10], surface slicks may be an important habitat feature for the neuston. Without dense aggregations of surface life, it is hard to imagine how these species would survive.

We also found evidence that some neuston may reproduce in the NPGP: at least one sample contained many small *Velella* roughly 1/2 cm in total length, and *Janthina* sp. and *Porpita* sp. less than 1 mm in length. While no direct growth rates have been measured for any neustonic organism, Bieri [33] inferred growth rates for *Velella* based on the size of stranded animals over time. Using these estimates, our sampled *Velella* would be only 7-23 days old. That is not sufficient time to drift far, suggesting that *Velella, Janthina,* and *Porpita* may be reproducing in the NPGP. This does not exclude the possibility that species are also being transported in or out of the NPGP. Several species of neuston, such as *Velella,* have remarkable windharnessing sails and can be dispersed long distances, and strand regularly along the US and Canadian west coasts [34]. Future studies should examine neuston growth rates, seasonal size and abundance, and the influence of wind on the distribution of neuston of different sizes.

If there is a neuston sea within the Garbage Patch, it may be an important resource for other species in the region. The endosymbiotic zooxanthellae of *Velella* and *Porpita* may allow them to persist even when the abundance of their prey (plankton) is low. In turn they may also be a more reliable food source for predators. Neuston are present in the diet of Pacific sea turtles [35,36], and Laysan albatross [37]. Juvenile loggerhead turtles have been caught in the region with rafting surface life in their stomachs [38]. Predation on neuston may also explain why these animals are sadly infamous for mistakenly ingesting large amounts of plastic.

In the North Atlantic Sargasso Sea, neuston provide a nursery ground, feeding ground, and habitat [15]. Similar to the Sargasso Sea, our results suggest the central NPGP has high surface life densities relative to surrounding waters, and this opens the interesting possibility that it may also serve similar ecological functions to the North Atlantic Sargasso Sea. Like in the Sargasso Sea, high densities of neuston in the NPGP may be due to both physical factors that concentrate neuston in this region, such as currents and wind, and biological factors such as temperature tolerance and life cycle characteristics.

Our findings suggest subtropical gyres and other areas of high plastic concentration may be more than just garbage patches. The presence of unnoticed neuston seas not only has implications for regional ecology and ecosystem services but possibly for international policy and biodiversity protection. The United Nations is currently negotiating a treaty to protect high seas biodiversity [39], and the Sargasso Sea has been identified as a key area of conservation interest [40]. If the Sargasso Sea is in fact only one of several neustonic seas, these other regions are worthy of future study and special consideration. Integration of observing systems of marine debris and biodiversity [41] would advance our understanding of the complex dynamics and multi-disciplinary interactions in the pelagic ocean. Ecological studies on the food webs and life history of neustonic species will allow us to better understand their temporal cycles and connectivity, and community science will help us better map life at the air-sea interface.

## Acknowledgments

Acknowledgments We thank Ben Lecomte and the crew of the Vortex Swim for generously providing us with an opportunity to collect samples, and Dr. Sara-Jeanne Royer for providing the Manta trawl. We would like to thank the organizers and attendees of the “The Ocean Cleanup Symposium 2019” at the University of Liverpool Institute for Risk and Uncertainty. This study is partly supported by NASA through Grant 80NSSC21K0857, funding the GO-SEA project (www.goseascience.org). NM and JH are also partly supported by NASA grants 80NSSC17K0559 and NNX17AH43G.

## Notes

### Competing Interest Statement

The authors have declared no competing interest.

### Summary of Updates

updated citations; genus names italicized in supplemental figures; correction of small typographical errors in the text.

