## Supplementary material for "High Concentrations of floating life in the North Pacific Garbage Patch": Chong_supplemental_figures_20May2022.docx

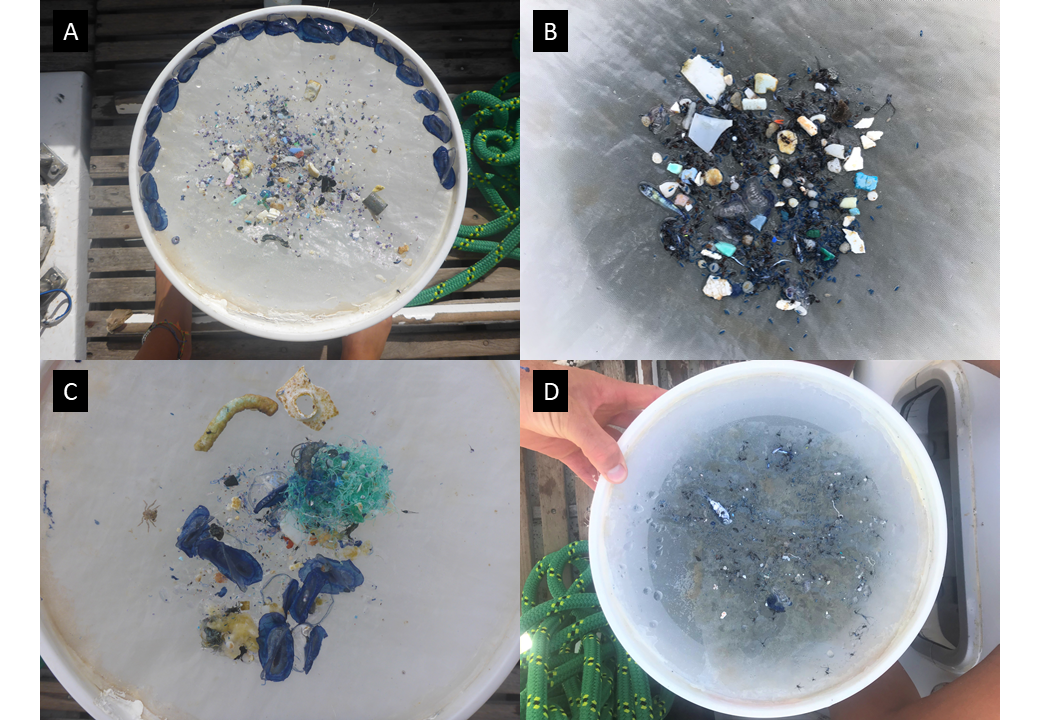

**Figure S1**. Examples of images taken by The Vortex Swim expedition team, showing contents of four neuston net / manta trawls. In all photos there is a mixture of neuston life and plastic fragments. *Velella* individuals can be clearly seen in A and C. *Halobates* and copepods are common in B. D is dominated by some gelatinous mass with not much plastic or true neuston. A. TLS_138, B. SJR_007, C. SJR_38, D. TLS_101.

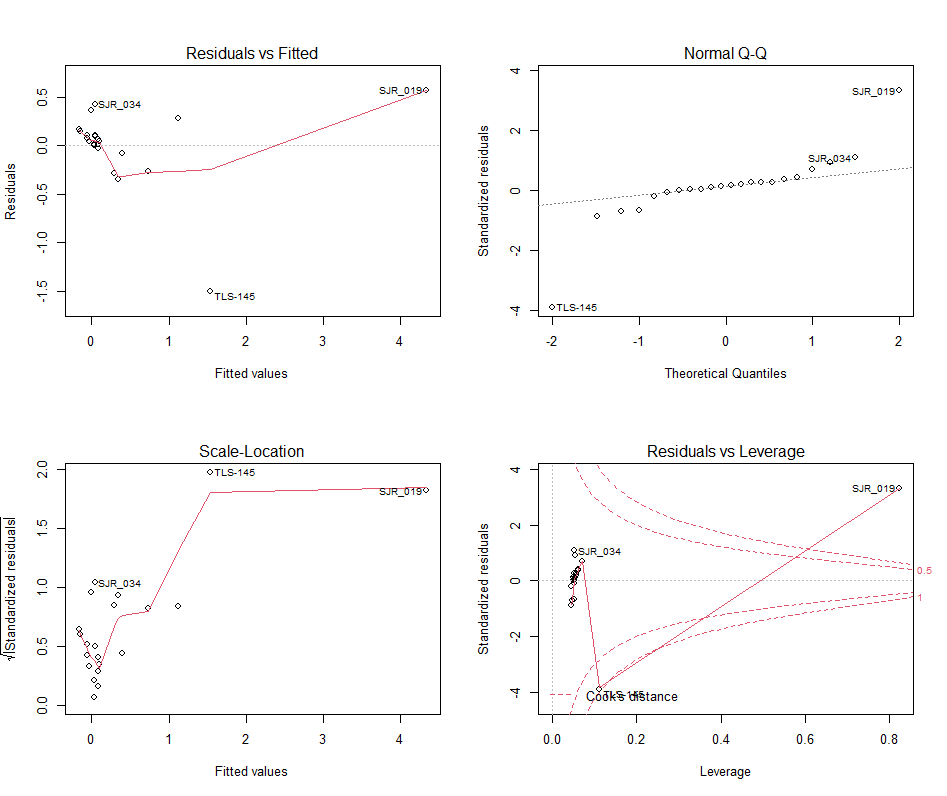

**Figure S2.** Regression diagnostics for the linear regression of plastic and true neuston densities. Bottom right: Residuals vs Leverage plot shows SJR_019 to be an influential sample with Cook’s distance > 1.

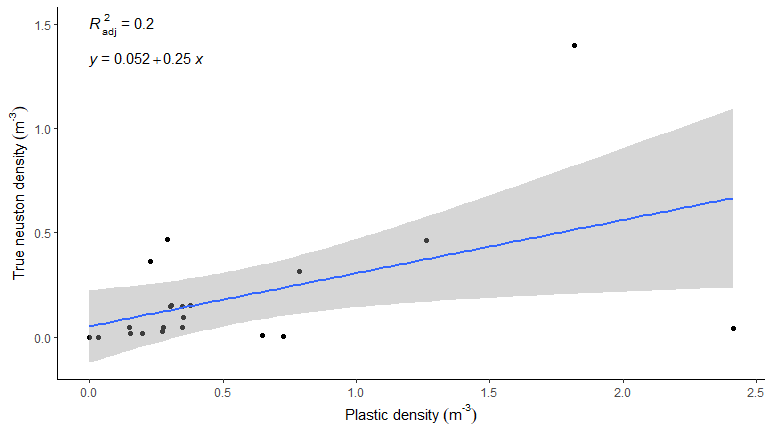

**Figure S3.** Linear regression with the outlier observation SJR_019 removed. Black points represent sites visited by the Vortex Swim expedition. Blue lines: linear regression. Grey regions show the 95% confidence band for each model. Regression line equations and adjusted R^2^ values are also displayed.

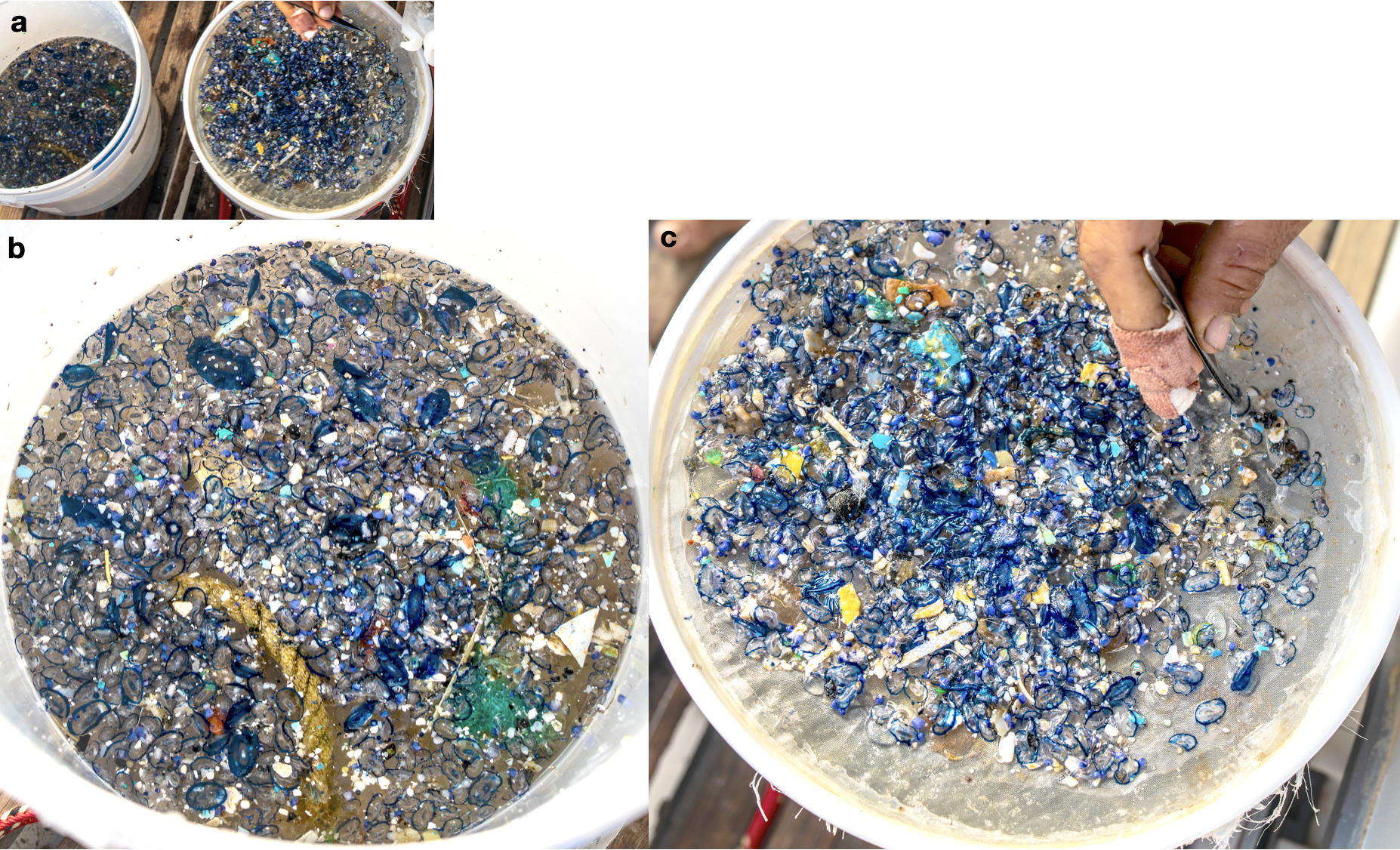

**Figure S4.** High concentrations of plastic and neuston from a sea surface slick (sample SJR_19). a. The full sample, in two batches. b. half the sample is in a bucket and c. the other in the sieve. Visual key: *Velella* are visible as blue rings, *Janthina* are smaller and appear violet and spherical, *Porpita* resemble blue bottoms with a central white disk. *Glaucus* are also visible.
