## Supplementary material for "High Concentrations of floating life in the North Pacific Garbage Patch": VS.R

```
rm(list = ls())
library("ggplot2")
library("ggpubr")
library("gridExtra")
library("dplyr")


#import csv file
averages_no_rounding <- read.csv("averages_no_rounding.csv")


#to subset the dataset
df <- select(averages_no_rounding, `Sample.name`, `Visit.order`, `Patch`, TN.vol, `plastic.vol`, `model.tracer.conc`)
#rename headings
names(df)[4] <- "true_neuston"
names(df)[5] <- "plastic"
names(df)[6] <- "tracer"
rownames(df) <- df[,1]
  
#lm of plastic conc and neuston conc: Figure 4
  lm_plastic_neuston <- ggplot(df, aes(plastic, true_neuston)) + geom_point() + geom_smooth(method = lm) +
    labs(y= bquote('True neuston density'~(m^"-3")), x = bquote('Plastic density'~(m^"-3"))) + theme_classic()+
    stat_regline_equation() + stat_regline_equation(label.y = 3.8, aes(label = ..adj.rr.label..))
  model1 <- lm(true_neuston~plastic, data = df)
  print(summary(model1))
  plot(lm_plastic_neuston)
#diagnostics
par(mfrow = c(2,2))
plot(model1) # diagnostics: SJR_019 had Cook's distance > 1; Figure S2
  
#lm of plastic conc and neuston conc without SJR_019: Figure S3
  lm_plastic_neuston_outlier_removed <- ggplot(subset(df,Sample.name!="SJR_019"), aes(plastic, true_neuston)) + geom_point() + geom_smooth(method = lm) + 
    labs(y= bquote('True neuston density'~(m^"-3")), x = bquote('Plastic density'~(m^"-3"))) + theme_classic()+
    stat_regline_equation() + stat_regline_equation(label.y = 1.5, aes(label = ..adj.rr.label..)) 
  model2 <- lm(true_neuston~plastic, data = subset(df,Sample.name!="SJR_019"))
  print(summary(model2))
  plot(lm_plastic_neuston_outlier_removed)
  #diagnostics
  par(mfrow = c(2,2))
  plot(model2)
  
# cube root transformation
  df$cubert_true_neuston <- df$true_neuston^(1/3)
  df$cubert_plastic <- df$plastic^(1/3)
  
  lm_plastic_neuston_cubert <- ggplot(df, aes(cubert_plastic, cubert_true_neuston)) + geom_point() + geom_smooth(method = lm) + 
    labs(y= bquote('cube root True neuston density'~(m^"-3")), x = bquote('cube root Plastic density'~(m^"-3"))) + theme_classic()+
    stat_regline_equation() + stat_regline_equation(label.y = 1.5, aes(label = ..adj.rr.label..)) 
  plot(lm_plastic_neuston_cubert)
  modelcubert <- lm(cubert_true_neuston~cubert_plastic, data = df)
  print(summary(modelcubert))
  par(mfrow = c(2,2))
  plot(modelcubert)

# boxplots for in and out of patch, KW tests-------------------------------

  #rearrange dataframe
  df2 <- data.frame(concentration = c(df[,"true_neuston"], df[,"plastic"]))
  df2$Patch <- c(df[,"Patch"], df[,"Patch"])
  df2$Sample.name <- c(df[,"Sample.name"], df[,"Sample.name"])
  df2$tracer <- c(df[,"tracer"], df[,"tracer"])
  df2$type[1:22] <- "True neuston"
  df2$type[23:44] <- "Plastic"

#boxplots to show plastic and true neuston in and out of patch: Figure 3  
p <- ggplot(df2, aes(x=Patch, y=concentration, fill=type)) + 
    geom_boxplot()+
    labs(x="In or Out of the NPGP", y = bquote('Particle density'~(m^"-3"))) +
    scale_color_grey() + theme_classic()
plot(p)

#KW test for plastic or neuston in and out of patch, find median and standard error
kruskal.test(true_neuston~Patch, data = df)
kruskal.test(plastic~Patch, data = df)

neustonin <- df$true_neuston[6:17]
neustonout <- df$true_neuston[c(1:5,18:22)]
plasticin <- df$plastic[6:17]
plasticout <- df$plastic[c(1:5,18:22)]

median(neustonin)
IQR(neustonin)

median(neustonout)
IQR(neustonout)

median(plasticin)
IQR(plasticin)

median(plasticout)
IQR(plasticout)
```
